# Early HIV-1 Gag Assembly on Lipid Membrane with vRNA

**DOI:** 10.1101/2023.01.27.525415

**Authors:** Anne X.-Z. Zhou, John A. Hammond, Kai Sheng, David P. Millar, James R. Williamson

## Abstract

Mass photometry (MP) was used to investigate the assembly of myristoylated full-length HIV-1 Gag (myr-Gag) and vRNA 5’ UTR fragment in a supported lipid bilayer (SLB) model system. The MP trajectories demonstrated that Gag trimerization on the membrane is a key step of early Gag assembly in the presence of vRNA. Growth of myr-Gag oligomers requires vRNA, occuring by addition of 1 or 2 monomers at a time from solution. These data support a model where formation of the Gag hexamers characteristic of the immature capsid lattice occurs by a gradual edge expansion, following a trimeric nucleation event. These dynamic single molecule data involving protein, RNA, and lipid components together, provide novel and fundamental insights into the initiation of virus capsid assembly.

## Introduction

The global impact of AIDS has made human immunodeficiency virus (HIV) one of the most intensely studied viruses. To infect new cells, HIV must assemble infectious viral particles within the infected host. This assembly process is complicated and carefully choreographed among viral proteins, virus genomic RNA (gRNA), and the inner leaflet of the plasma membrane, involving many cellular host proteins.^1^ The viral structural polyprotein Gag, accounting for more than half of the virus biomass, plays a crucial role in this process, with involvement of four domains with different but coordinated functions: the matrix (MA) domain mediates binding to the membrane and specifically interacts with phosphatidylinositol 4,5-biphosphate (PI(4,5)P_2_) on the membrane, the capsid (CA) domain mediates association with other Gag proteins, the nucleocapsid (NC) domain confers selective binding to gRNA, and the p6 domain is responsible for recruiting cellular host factors.^2^ The general functions of Gag are to oligomerize, selectively bind viral gRNA in dimeric form, and interact with the inner leaflet of the plasma membrane to form newly assembled immature virus particles.

Although virus assembly has been studied extensively over the years,^3–7^ there are only a few studies involving the three key components: myristoylated full-length Gag, viral RNA, and lipids. Recent studies using two of these three components have implicated the intertwined interactions between Gag, gRNA, and lipids. Recent structural and biochemical results have identified gRNA dimer can facilitate Gag-RNA nucleation,^8^ while there is mounting evidence showing that Gag-Gag interactions, via both CA domain and MA domain, are the main driving forces for the final immature lattice formation.^9,10^ Within the Gag protein, the CA domain forms stable hexamers in the immature lattice, and the Gag MA domain forms a hexamer of trimers on a model membrane system.^11–13^ The two domains are connected by a flexible linker, possibly allowing both domains to adopt their preferred oligomerization pattern at the same time.^11,13^ The interactions between MA domain and the lipid membrane add yet another layer of complexity to the process, and there is no consensus as to how oligomeric Gag complexes grow and where those additional Gag proteins come from. Thus, it is essential to investigate this important viral assembly process with all three components present in the system.

To monitor engagement of Gag on the membrane with gRNA in a single molecule approach, we harnessed the power of the newly developed technique of mass photometry. Mass photometry is based on principles of interferometric scattering microscopy (iSCAT) microscopy, where the minute amount of light scattered by single molecules near a slide surface can be detected and transformed into a contrast by applying a model point spread function (PSF), and after calibration, be converted to molecular weight in a label-free manner.^14,15^ Such correlation between the amount of light scattered by a particle and the molecular weight of the particle is linear throughout a wide range, regardless of shape and species. Mass photometry data can be acquired in two observation modes. In a conventional mass photometry assay (landing assay), one can detect interferences of individual molecules as they adsorb nonspecifically on the slide surface, which then can be converted to a molecular weight by calibration with a set of known standards. By observing a sufficiently large number of adsorption events, a molecular weight distribution of species present in solution can be developed. In addition, with a different background subtraction algorithm, a mass photometer instrument can also perform a mass-sensitive particle tracking (MSPT) assay to track individual molecules diffusing on a surface. In this mode, the changes in contrast patterns along the positional trajectory are recorded, allowing both positional dynamics and changes in the molecular weight to be observed directly at the same time (**Fig. 1, S1**).^16,17^

**Fig. 1.**
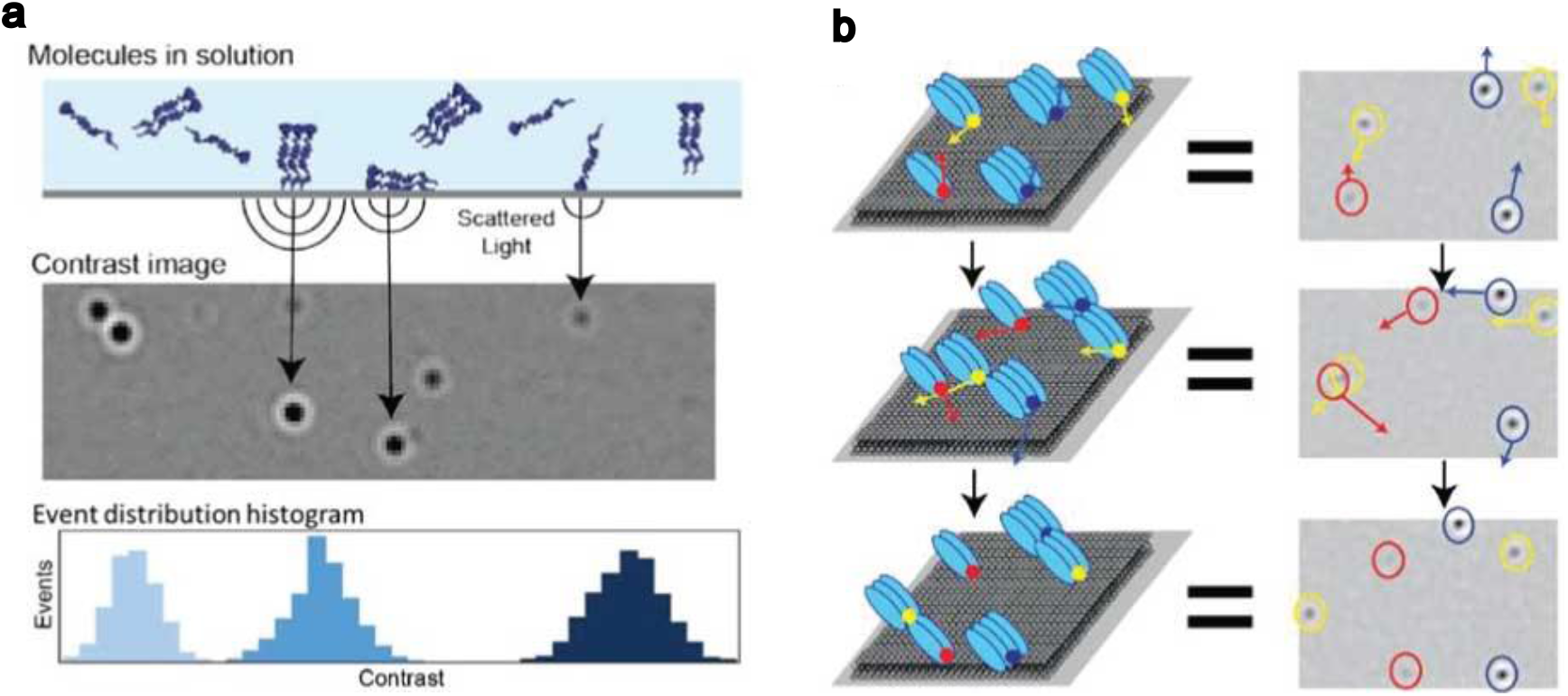
Schematic for mass photometry. (**a)** Schematic for mass photometry landing assay, for determination of the distribution of molecular weights in solution. As individual molecules or complexes diffusing in solution absorb nonspecifically to the glass surface, interference of scattered light gives rise to contrast features over time. The contrast is proportional to molecular weight, and a histogram of contrast events gives the distribution of species in solution. (**b)** Schematic for MSPT where oligomeric complexes are diffusing in a 2D membrane. The position and contrast (which can then be converted to molecular weight) are recorded as trajectories in time.

Using mass photometry, we examined the early steps in Gag assembly using myristoylated full-length Gag (myr-Gag), 5’ UTR gRNA containing the packaging signal,^18^ and a supported lipid bilayer (SLB) containing PI(4,5)P2. We demonstrated the importance of the presence of lipid and gRNA in Gag oligomerization and showed that proper CA-mediated Gag dimerization is needed for assembly and efficient recruitment of gRNA to the membrane. Our data indicate that an MA-mediated trimer may be the smallest nucleating unit on the membrane, and that the assembly proceeds via an edge-expansion process, where one or two Gag proteins are added at a time from the solution to the growing assembly nucleus.

### Myr-Gag primarily consists monomers and dimers in solution

The MP landing assay was used to monitor the molecular weight distribution of myr-Gag or myr-Gag-RNA complexes in binding buffer. Over the concentration range tested (10-100 nM), myr-Gag in the binding buffer always has a mass distribution consisting of a ~55 kDa peak and a ~110 kDa peak, corresponding to a myr-Gag monomer and dimer, respectively (**Fig. S2a**) When adding WT 5’ UTR RNA (352 nt, 110 kDa) together with 50 nM myr-Gag, an additional small peak was observed, corresponding to an RNA dimer (220 kDa) (**Fig. S2b**). The 5’-UTR RNA used here is well known to form a dimeric species,^19,20^ which was confirmed using non-denaturing gel electrophoresis. Although a few particles with higher molecular weights were detected, the distribution of lower masses suggests that Gag-RNA complexes are not significantly populated in solution (**Fig. S2b**). Neither myr-Gag alone nor myr-Gag with RNA form significant higher order complexes in solution.

### HIV-1 5’ UTR RNA promotes further Gag assembly on SLBs

MSPT experiments were employed to track myr-Gag diffusion on a lipid membrane. Diffusing complexes were readily observed and tracking of individual complexes yielded many trajectories suitable for analysis (**Fig. S1**). The myr-Gag concentration and the PI(4,5)P2 composition in the SLB were surveyed to identify 50 nM myr-Gag and 2% PI(4,5)P2 as a condition yielding a reasonable trajectory density within the field of view (FOV) that optimizes the quantity and quality of the data that can be extracted (**Fig. S1e**).

To test whether Gag-RNA complexes form on an SLB and whether RNA promotes Gag assembly on the membrane, myr-Gag was examined in MSPT experiments on an SLB, alone and with varying amounts of WT 5’ UTR RNA. The results show a remarkably rich set of complexes forming in the presence of RNA, with an informative concentration dependence (**Fig. 2**). In the absence of RNA, a major peak corresponding to a myr-Gag trimer was observed, in contrast to the monomer/dimer species that predominate in solution. This observation strongly suggests that Gag-Gag interactions are mediated by Gag domains proximal to the SLB, presumably the MA domain. Significant Gag-RNA binding is observed at a very low RNA concentration (0.1 nM), with the appearance of a peak at ~385 kDa, corresponding to an RNA_2_:Gag_3_ complex (**Fig. 2**). Small populations of larger complexes also appear, corresponding to higher order oligomeric states of Gag (**Fig. 2**). However, the myr-Gag trimer peak remains as the dominant species. As the RNA concentration was increased further up to 2 nM, the myr-Gag trimer peak gradually disappears as a broad peak representing RNA_2_:Gag_3/4/5_ increases, together with a significant distribution of other higher-MW complexes (n.b. Gag-RNA complex peak assignment rationale is described in Methods).

**Fig. 2.**
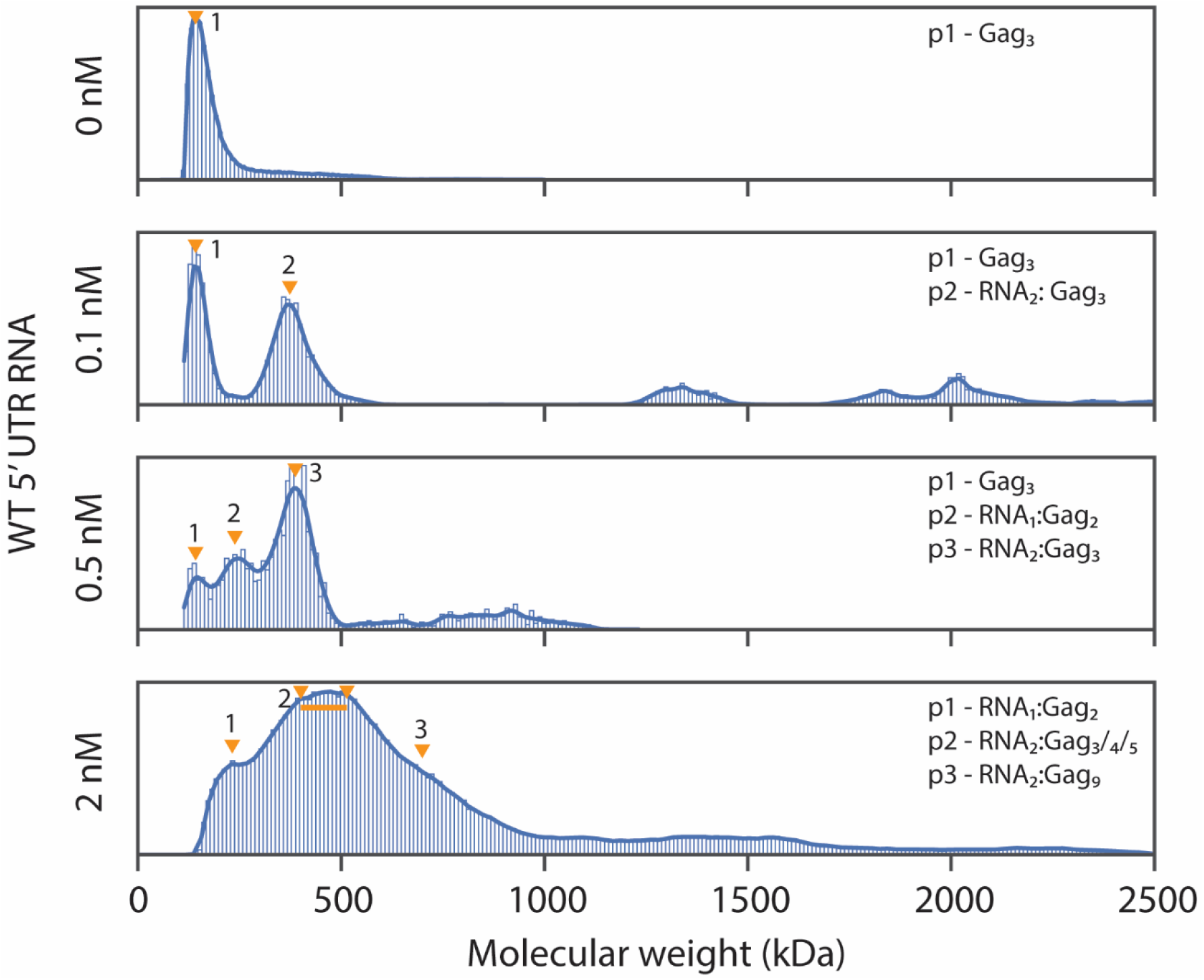
Molecular weight distribution of Gag with 5’ UTR RNA. Molecular weight distribution probability density plot of 50 nM myr-Gag mixed with different concentrations of WT 5’ UTR RNA on SLB. Labeled complex composition were estimated from kernel density plot peak molecular weights. All sample sizes are larger than 2500 and collected from at least 3 different recordings.

These results demonstrate that the presence of even minute amounts of 5’ UTR RNA enhances Gag oligomerization on the membrane, and that not only RNA concentration but also the protein-to-RNA ratio affects the final distribution of Gag-complex species on the SLB. In summary, myr-Gag preferentially forms dimers in solution without and with 5’ UTR RNA, but forms trimers or multimers when binding to dimeric 5’ UTR RNA in the context of an SLB. This is clear evidence for the cooperative interaction between protein, RNA, and lipid in the earliest steps of capsid assembly.

### Gag dimerization is important for Gag assembly on a SLB

Gag forms dimers through interactions between the capsid domain (CA), and CA-CA interactions are a key feature of the 2-fold interactions in the Gag immature lattice.^10^ To investigate the effect of Gag CA dimerization in the assembly process on the membrane, the MWM myr -Gag mutant (M39A, W184A and M185A disrupting CA-CA dimer interface) was expressed, purified, and tested in mass photometry experiments.^21^ In solution, MWM myr -Gag forms a smaller proportion of dimers compared to WT myr-Gag, as expected (**Fig. 3**). In the SLB, MWM myr-Gag also formed Gag_3_ as the predominant species (**Fig. 3**). Gag forms trimers through interactions between the MA domains, which is a key feature of the 3-fold interactions in the Gag immature lattice.^22–24^ Formation of Gag trimers in the MWM myr-Gag in the context of SLBs is consistent with an MA-mediated oligomerization that is unaffected by the CA dimerization mutations. The mass distribution in the supernatant after SLB incubation was also measured in a landing assay, to determine if the distribution of species in solution was altered by exposure to the SLB. In both cases, there was one broad peak between 55 kDa and 110 kDa, indicating no species larger than a Gag dimer exists in the solution after SLB incubation (**Fig. 3**).

**Fig. 3.**
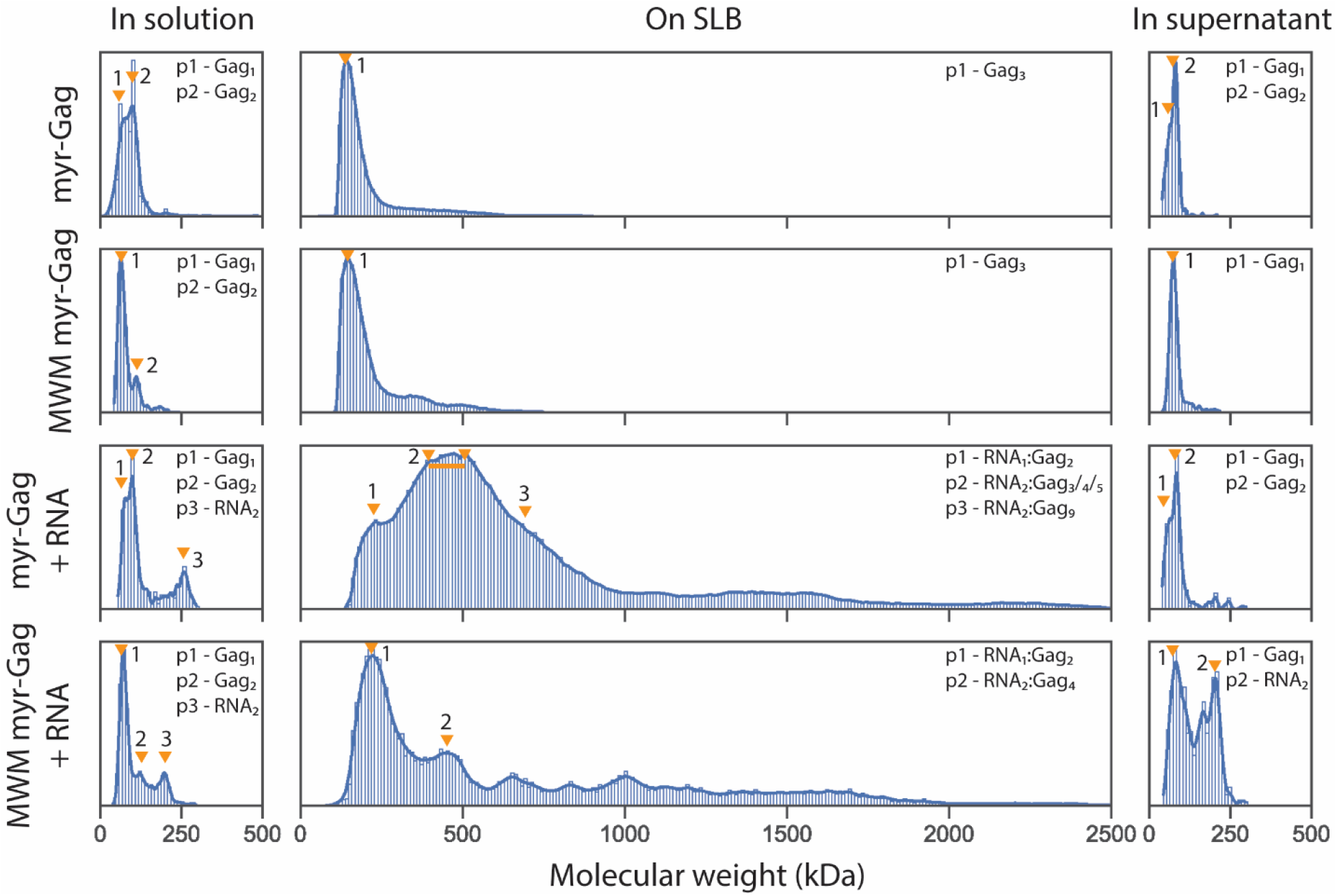
Molecular weight distribution of WT myr-Gag and MWM myr-Gag mutant with 5’ UTR RNA. Molecular weight distribution probability density plot of 50 nM WT myr-Gag or MWM myr-Gag without or with 2 nM 5’ UTR RNA in binding buffer before applying to the SLB (landing assay), on SLB (MSPT assay), or in supernatant solution after SLB incubation (landing assay). Labeled complex composition were estimated from kernel density plot peak molecular weights. All sample sizes are larger than 2500 for MSPT assay data and larger than 400 for landing assay data. Data were collected from at least 3 different recordings.

When RNA is added to the mixture, MWM myr-Gag does not bind to RNA significantly in solution, similar to WT myr-Gag (**Fig. 3**). However, when applied to the SLB surface, the mutant showed fewer higher order complexes and only a small fraction of Gag_4_:RNA_2_, compared to myr-Gag (**Fig. 3**). Also, the MWM mutation did not abolish the RNA_1_:Gag_2_ complex formation (~220 kDa) as expected based on the mutation sites, indicating this complex can be maintained via either Gag-RNA interactions or Gag MA-MA interactions in the context of the SLB. It is important to note that while MSPT data can discern the molecular weight of complexes, it cannot show that the geometry of a given complex is homogeneous for a given sample, or the same for different samples. The composition of the supernatant after SLB incubation also showed different distributions. The MWM myr-Gag complexes exhibited a much larger portion of RNA dimer after 30 min SLB incubation compared to the WT Gag, indicating that MWM myr-Gag cannot recruit 5’ UTR RNA to the membrane surface as efficiently as the WT myr-Gag (**Fig. 3**).

### Gag assembly occurs in small incremental steps

Step detection on time-resolved mass traces was done using the Kalafut-Visscher algorithm without manual step-size input,^25^ allowing detection of transitions between species on the SLB. Comparing the results of myr-Gag alone and myr-Gag with WT 5’ UTR RNA, the step size distributions for attachment events are largely similar, with the myr-Gag with WT 5’ UTR RNA group having more large steps (**Fig. 4a**). Both groups have the smallest, and also the most abundant, step size peak at the molecular weight of either 1 or 2 myr-Gag (~55 and 110 kDa), signifying that Gag assembly occurs primarily by adding one or two Gag proteins at a time (**Fig. 4a**). In addition, the transition density histogram showed that this step distribution is not limited to formation of the very first RNA:protein nucleus, and addition of 1 or 2 myr-Gags was the major transition observed for complexes of all sizes detected (**Fig. 4c, 4d**).

**Fig. 4.**
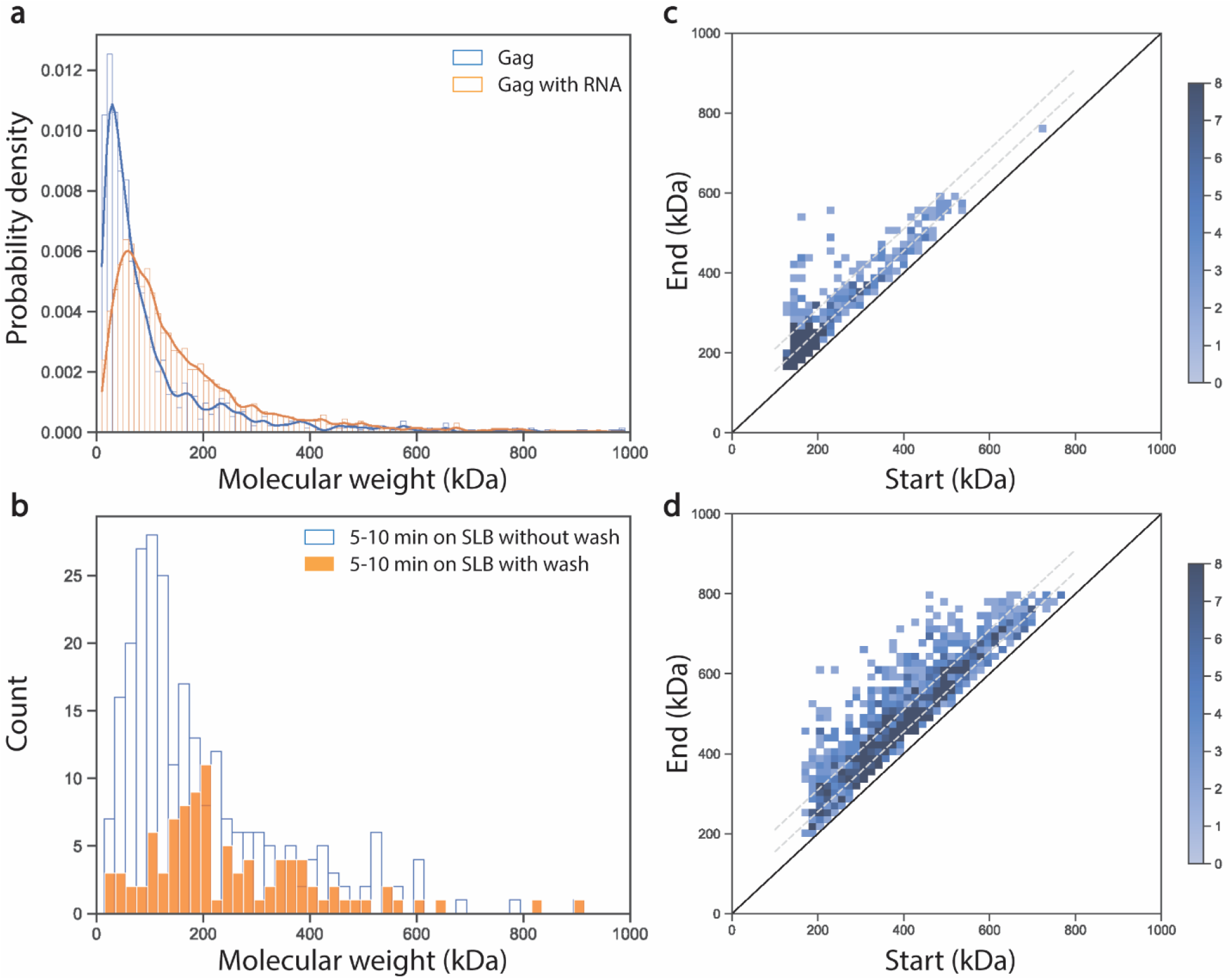
Step size distribution of attachment events and step analysis. (**a)** Probability density of step size distribution for attachment events for 50 nM myr-Gag on SLB (blue) and 50 nM myr-Gag with 2 nM WT 5’ UTR RNA on SLB (orange). (**b)** Histogram of step size distribution for attachment events of 50 nM myr-Gag with WT 5’UTR RNA at 5-10 min after incubation on SLB (blue) and 50 nM myr-Gag with 2 nM 5’ UTR RNA at 5-10 min after incubation on SLB and buffer wash at 5 min (orange). (**c)** Transition density plot of step size distribution for attachment events for 50 nM myr-Gag on SLB, with the two dotted grey line shows the “+ 55 kDa” steps and “+ 110 kDa” steps. (**d)** Transition density plot of step size distribution for attachment events for 50 nM myr-Gag with 2 nM 5’ UTR RNA on SLB, with the two dotted grey line shows the “+ 55 kDa” steps and “+ 110 kDa” steps.

Considering the results of the landing assay which showed Gag in solution does not form larger complexes before or after SLB incubation (**Fig. 3, Fig. S2b**), it is likely that Gag directly adds to the Gag complex cores on the membrane from the solution, instead of from lateral collision on the membrane during the early stages of Gag assembly on the SLB. When RNA is present, the chances of adding two or more Gag’s at a time is higher, possibly because RNA provides an assembly scaffold. Indeed, there were no observed events that appeared to correspond to collisions between complexes within the SLB resulting in a higher molecular weight species. To test this hypothesis, the supernatant containing Gag and RNA was washed away after 5 min incubation on the SLB, and the step size distribution for attachment events in the next 5 min were recorded and compared to a control group without wash at the same time frame. The result showed that there is a sharp decrease in the most abundant Gag or Gag_2_ steps when the Gag protein reservoir in solution was removed, while RNA_1_:Gag_2_ attachment events remained, supporting the hypothesis that most attachment events are coming from the solution (**Fig. 4b**).

The combination of mass photometry landing assay and the MSPT assay has led to new insights into the early stages of the Gag assembly process. With the data discussed above, we showed the importance of incorporating all three components (myristoylated full-length Gag, 5’ UTR RNA, and the lipid membrane) in the experimental system (**Fig. 2**). We also demonstrated the different Gag complex distributions in solution and on membrane, with Gag monomers/dimers dominant in solution, and Gag trimers dominant on the SLB (**Fig. 2, S2**). Mutant Gag abolishing CA dimerization showed a defect in forming early Gag complex species and a compromised RNA recruitment ability (**Fig. 3**). Attachment step analysis also revealed that Gag in solution directly adds to the assembly core on the SLB by steps of one or two Gag proteins at a time (**Fig. 4**).

While a Gag hexamer is the smallest repeating structural unit in the immature Gag lattice, there has been no consensus on the potential role of a Gag hexamer as an assembly intermediate, despite decades of studies.^26^ Our data do not support a Gag hexamer being a major assembly step, as we did not observe steps containing six Gag proteins. There has been evidence showing Gag adds to the immature lattice via lower order multimers,^27^ and more recently, patterns of an edge expansion model were identified in the immature lattice of a budded virion and on mica surface in AFM experiments,^28,29^ but no similar study has been done on a biologically relevant membrane system focusing on the early assembly steps. Our data support an edge expansion model in early-stage Gag assembly where a Gag hexamer is neither the nucleus for assembly, nor added as a whole to the nucleus. Rather, assembly proceeds by addition of Gag monomers and dimers to the edge of an assembly core on a lipid membrane surface where trimerization, mostly mediated by the MA domain, plays a crucial role.

The interplay between Gag monomers and dimers in solution with Gag trimers in the membrane may reflect the duality of the immature lattice when viewed from the standpoint of the MA and CA domains. Lattice models exist for hexamers-of-trimers for the MA domain (**Fig 5a**), and trimers-of-hexamers for the CA domain (**Fig 5b**), that have similar spacings and 2-fold, 3-fold, and 6-fold axes. However, cryo-electron tomographic analysis of immature virus particles indicates that the MA and CA lattices can be independently identified, but do not order and superimpose directly, presumably due to the flexible linker between the MA and CA domains.^23^ Several molecular models have been developed by superposition of the symmetry axes of the MA and CA lattices,^11,13^ which can be illustrated schematically in **Fig. 5c**. There is a key feature of such a model that has important implications for the observed data and resulting assembly model: any given Gag involved in a trimer via MA-MA interactions does not have a common oligomeric partner with any Gag also involved in its hexamer via CA-CA interactions (**Fig. 5a**). In effect, satisfying the symmetry of both MA and CA lattices may in fact limit the addition of oligomers to the nucleus to one or two at a time.

**Fig. 5.**
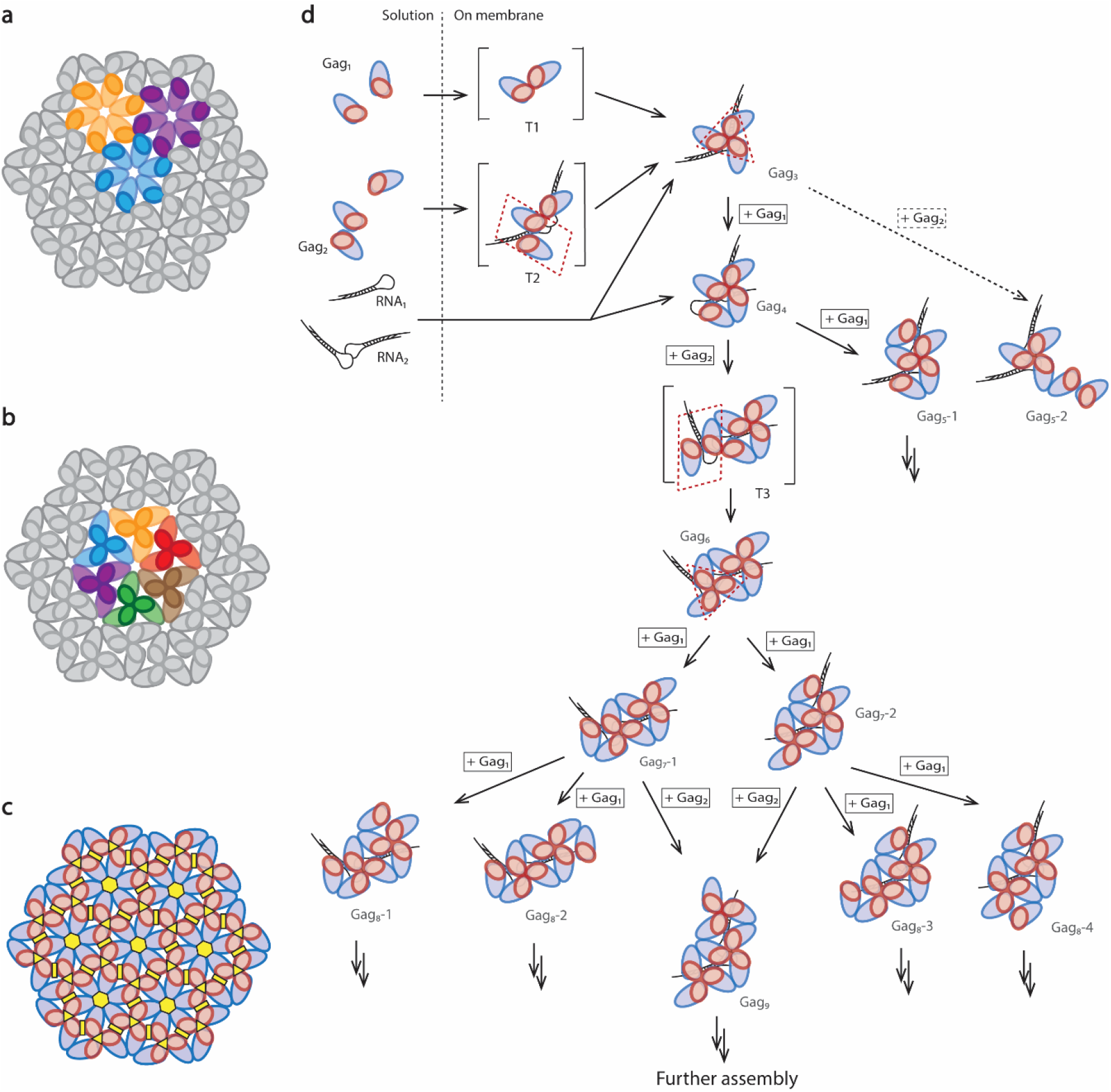
Schematics for Gag lattice and Gag assembly. (**a)** Schematic of a trimer of Gag hexamers. (**b)** Schematic of a hexamer of Gag trimers. (**c)** Schematic of Gag lattice with symmetry axis. Gag MA domain is represented in red and Gag CA domain is represented in blue, with six-fold symmetry axis labeled in yellow. (**d)** Schematic of proposed assembly pathway for early steps. Gag docks first on the membrane to form the initial Gag_3_ assembly core, driven by the trimer interface of both MA and CA. Additional Gag monomers and dimers then add to the core directly from the solution. When a Gag dimer (CA-mediated) is added in one step, it is proposed here that the dimer would first dock via an incomplete trimeric surface (MA-mediated, noted in square brackets) and then quickly shift its conformation to complete that MA-mediated trimeric interface. Gag monomers and dimers then add to the complex in an orderly pattern as exemplified above, gradually forming complete hexamers. There can be other alternative pathways expanding the complex toward different directions (possible pathways noted by arrows and less possible pathway noted by a doted arrow, further assembly noted by double arrows), but attachment events must be a combination of Gag monomers and dimers to complete all dimer and trimer interfaces in each step. Step sizes are noted in squares and possible isoforms of Gag complexes are noted in dashed numbers.

Considering the complex orchestration of dimeric interactions, trimeric interactions, and hexametric interactions between Gag proteins, and based on the observed complex molecular weight distributions and step size distributions for attachment events, a detailed assembly path is proposed here (**Fig. 5d**). As Gag monomer and Gag dimer are the only large populations in solution, we propose that they will dock first on the membrane to form a transition complex T1 or T2, both containing an incomplete MA trimer interface with T2 also harboring a completed CA dimer interface. Such transition complex would then quickly rearrange into Gag_3_, driven by the trimer interface of both MA and CA. This initial step is then followed by addition of another Gag monomer, forming a Gag_4_ complex with a complete trimer interface and a dimer interface, satisfying both MA interaction and CA interaction. Next, a Gag dimer can add to the Gag_4_ complex to form another transition complex T3, which, similar to T2, would also quickly rearrange into Gag6 to complete another MA trimer interface to form Gag_4_. While a partial hexamer can be relatively stable,^28^ the fact that Gag prefers trimer over dimer as the smallest complex on SLB suggests that an exposed dimer interface is more tolerable than an incomplete trimer interface, limiting further assembly pathway via Gag5-2. Gag monomers and dimers then add to the complex in an orderly pattern as exemplified above, gradually forming complete hexamers. Since the MWM myr-Gag mutant was shown to significantly disrupt RNA recruitment (**Fig. 3**) and RNA_1_:Gag_2_ may add to the assembly core via diffusion on the membrane (**Fig. 4b**), the incorporation of RNA into the early assembly core is very likely to happen together with Gag dimer recruitment. There can be other alternative pathways expanding the complex toward different directions (dotted arrows in **Fig. 5**), but attachment events have to be a combination of Gag monomers and dimers to complete all dimer and trimer interfaces in each step. When Gag dimerization was compromised by the MWM mutation, the formation of transition complexes is also largely impaired, resulting in fewer Gag_4_ complexes and almost no larger complexes, signifying that adding Gag monomers alone would not sustain the edge expansion.

Our study demonstrated the potential of mass photometry in studies involving protein-protein assembly on a membrane surface. This is the first time that Gag MA domain trimerization has been considered as an important driving force of early-stage Gag assembly on the membrane, instead of Gag CA domain hexamerization. It is also the first study to support an edge expansion model of myristoylated full-length Gag during early stages of assembly on a lipid membrane. The dominant role of MA in forming the nucleus may be a consequence of its proximity to the membrane via the myristoyl anchor. Once sufficient clustering of Gag occurs by addition of monomers/dimers, the formation of CA hexamers is supported by edge expansion. Remarkably, the dual symmetry nature of the MA and CA lattices imposes a stepwise assembly on the membrane, possibly to ensure smooth formation of an extended lattice by a lengthy series of identical steps. This work integrates many known features of Gag assembly, but provides key insights into the earliest steps of forming the essential lipid-RNA-protein nucleus for RNA packaging and provirion assembly.

## Methods

### Materials

A complete list of plasmids and primers used in this work is provided in Tables S6 and S8. Primers and natural oligonucleotides were purchased from IDT (Coralville, Iowa). Sequencing was performed by Genewiz (San Diego, CA). Plasmids were purified using a commercial miniprep kit (D4013, Zymo Research; Irvine, CA). PCR products were purified using a commercial DNA purification kit (D4054, Zymo Research) and quantified by A260/A280 absorption using an Infinite M200 Pro plate reader (TECAN). All mass photometry data were collected on Refeyn Two^MP^, and data acquisition was done using Aqcuire^MP^. All experiments involving RNA species were done with RNase-free reagents, pipette tips, tubes and gloves to avoid contamination.

All DNA plasmid samples were stored at −20 °C. All RNA samples and protein samples were stored at −80 °C. All vesicles and nanodiscs were stored at 4 °C

### TEV expression and purification

TEV protease was expressed by transferring a plasmid encoding TEV under a CMV promoter to *E. coli*, with a TEV cleavage site inserted at the C-terminus of TEV and followed by a (His)_6_ affinity tag. Cell culture was grown at 37 °C until its OD_600_ reached 0.6, at which point IPTG (501964098, Fisher Scientific) was added to induce protein expression. Cells were collected 5 h after induction and stored at −80 °C before purification.

Cell pellets were resuspended in TEV sonication buffer (pH 8.0, 50 mM Na_2_HPO_4_, 200 mM NaCl, 10% glycerol, 1 mM β-mercaptoethanol) with 1 tablet of protease inhibitor and incubated on ice for 10 min. The mixture was then sonicated at 4 °C on ice (Qsonica Sonicator Q500 Sonicator, 20 s on, 59 s off, 50% amplitude, 8 min on in total). After sonication, 5% poly(ethyleneimine) (PEI) solution was gently added to the mixture dropwise while stirring at 4 °C to a final concentration of 0.1%, followed by centrifugation at 10,000 g for 30 min (Beckman coulter Allegra 25R centrifuge). The supernatant was removed and loaded onto a 5 mL HisTrap column (17-5255-01, Cytiva) equilibrated with TEV HisTrap A buffer (pH 8.0, 50 mM Na_2_HPO_4_, 200 mM NaCl, 10% glycerol, 25 mM imidazole). TEV was eluted from the HisTrap column by applying an increasing concentration gradient of HisTrap B buffer (pH 8.0, 50 mM Na_2_HPO_4_, 200 mM NaCl, 10% glycerol, 250 mM imidazole). Gradient fractions containing TEV were pooled together. EDTA and DTT were then added to the pooled fractions to final concentrations of 2 mM and 5 mM, respectively. The TEV fractions were then exchanged to TEV gel filtration buffer (pH 7.5, 15 mM Na_2_HPO_4_, 100 mM NaCl, 10% glycerol) and concentrated using a 10k concentrator. The mixture was then loaded onto a SEC200 size exclusion column (17-5174-01, Cytiva) equilibrated with TEV gel filtration buffer, followed by isocratic gel filtration buffer elution. Fractions containing TEV were pooled together and concentrated using a 10k concentrator. The final TEV solution was filtered through a 0.22 μm filter (SLGP033N, Millipore), flash frozen with liquid nitrogen, and stored at −80 °C in small aliquots. The presence of full-length TEV and its purity were monitored by SDS-PAGE.

### Gag expression and purification

A single plasmid was constructed bearing expression cassettes for both full-length Gag as a C-terminal MBP-(His)_6_ fusion, and the yeast NMT1 gene, and transformed into *E. coli* cells. The cell culture was grown at 37 °C until its OD_600_ reached 0.5 and myristic acid stock solution was added with a final concentration of 50 μM. The cell culture was then incubated at 28 °C for 15 min before adding 1 M IPTG to a final concentration of 0.5 mM. The incubation temperature was kept at 28 °C after induction, and the same amount of myristic acid was added to the culture again after 2.5 h incubation to maintain the myristic acid concentration before collecting the cells after 5 h incubation time after induction in total. The cell pellets were stored at −80 °C before purification.

The myristic acid solution was freshly prepared at 5 mM by first dissolving the myristic acid in 5 mL ethanol and then diluted with 50 mL pre-warmed 0.6 mM BSA at pH 9 (adjusted with NaOH). The solution remained at about 50 °C before use to maintain the solubility of myristic acid.

Cell pellets were resuspended in Gag sonication buffer (pH 7.5, 25 mM HEPES, 100 μM TCEP, 100 μM EDTA, 500 mM NaCl, 20 mM imidazole, 0.1% Tween 20) with 1 tablet of protease inhibitor and incubated on ice for 10 min. The mixture was then sonicated at 4 °C on ice (20 s on, 59 s off, 50% amplitude, 8 min on in total) and was then centrifuged at 10,000 g for 30 min. The supernatant was removed and loaded onto a 5 mL HisTrap column equilibrated with Gag HisTrap A buffer (pH 7.5, 25 mM HEPES, 100 μM TCEP, 100 μM EDTA, 500 mM NaCl, 20 mM imidazole, 0.1% Tween 20, 20 mM imidazole). Gag-MBP was then eluted from the HisTrap column by applying an increasing concentration gradient of Gag HisTrap B buffer (pH 7.5, 25 mM HEPES, 100 μM TCEP, 100 μM EDTA, 500 mM NaCl, 20 mM imidazole, 0.1% Tween 20, 400 mM imidazole) and fractions containing Gag-MBP were pooled together. The pooled fractions were then loaded onto a 5 mL Heparin column (17-0407-01, Cytiva) equilibrated with Gag Heparin A buffer (pH 7.5, 25 mM HEPES, 100 μM TCEP, 100 μM EDTA, 25 μM ZnCl_2_, 100 mM NaCl), followed by Gag Heparin B buffer (pH 7.5, 25 mM HEPES, 100 μM TCEP, 100 μM EDTA, 25 μM ZnCl_2_, 1 M NaCl) gradient elution. Fractions containing Gag-MBP were pooled together and concentrated using a 10k concentrator. The final Gag-MBP solution was exchanged into TEV cleavage buffer (pH 7.5, 50 mM Tris, 500 μM EDTA, 25 μM ZnCl_2_, 500 mM NaCl) and concentrated. TEV was added to the Gag-MBP solution, and the mixture was incubated at 30 °C for 2 h. The reaction was then loaded onto a Ni-NTA column (30410, QIAGEN) equilibrated with TEV cleavage buffer, followed by washing with 4 mL TEV cleavage buffer. All eluates were collected and concentrated at 15 °C to avoid Gag aggregation, as the solubility tag MBP had been removed. Glycerol was then added to the final Gag solution and the protein was stored at −80 °C in small aliquots. The presence of full-length Gag-MBP and its purity were monitored by SDS-PAGE and the molecular weight of the myr-Gag was confirmed by intact mass spectrometry.

To express Gag without myristoylation, a G2A mutation was introduced into the Gag sequence, which has been proven to abolish Gag myristoylation. All protein expression and purification procedures were the same aside from the protein encoding sequence.

### RNA preparation

Viral UTR RNA and its variants were made by *in vitro* transcription (IVT) (AmpliScibeTM T7-FlashTM, ASF3257, Lucigen) using a template containing a T7 promoter made from PCR off a plasmid encoding full-length gRNA sequence. IVT was done at 37 °C overnight, followed by spin-column purification. RNAs were stored at −80 °C in small aliquots.

Aliquots of RNA was thawed slowly on ice and then diluted to 1 μM in RNA refolding buffer (pH 7.5, 20 mM Tris, 10 mM NaCl, 140 mM KCl, 1 mM MgCl_2_). The diluted RNA was refolded by heating it up to 80 °C for 5 min followed by gradually cooling it down to 10 °C. The refolded RNA was kept on ice before use. The integrity of RNA was verified by urea SDS-PAGE and the conformation change through refolding was demonstrated by native agarose gel electrophoresis.

### SLB preparation

SLBs were prepared using defined mixtures of phosphatidyl choline (POPC) (850457, Avanti) and PI(4,5)P2 (840046X, Avanti) if needed. The lipids were dissolved in chloroform, mixed in the desired ratios, dried under nitrogen and the residual chloroform was removed in a vacuum desiccator overnight at room temperature. The dried lipid “cake” was then be resuspended in SLB buffer (pH 7.5, 20 mM Tris, 10 mM NaCl, 140 mM KCl). The lipid solution was then extruded through a polycarbonate membrane with a pore size of 50 nm using an Avanti mini-extruder (610020, Avanti) for 49 passes to form lipid vesicles with relatively uniform size. The SLBs were prepared by applying the solution of the vesicles to a cleaned glass slide surface immediately after plasma treatment (maximum power, 10 min at 0.5 bar air pressure). Unruptured vesicles were removed through extensive washing with SLB buffer after a 15-20 min incubation at room temperature to allow complete membrane formation.

### Mass photometry landing assay

Glass slides were washed by distilled water, isopropyl ethanol (34863, Sigma-Aldrich), water, isopropyl ethanol, and water again before being dried in an air stream. A gasket (3 mm diameter, 1 mm thickness, GBL103280, Grace Bio-Labs) was carefully placed on a cleaned slide, which was then placed on the mass photometer holder. Gag binding buffer (pH 7.5, 20 mM Tris, 10 mM NaCl, 140 mM KCl, 1 mM MgCl_2_, 50 μM ZnCl_2_) filtered by a 0.22 μm filter was added to the well. Sample protein solution or protein-RNA mixture was diluted or mixed in Gag binding buffer and incubated for 5 min before replacing the Gag binding buffer in the well, followed by immediate recording in the mass photometer of 1-2 min at a frame rate of 100 Hz.

### Mass photometry mass-sensitive particle tracking (MSPT) assay

Glass slides were washed with water, isopropyl ethanol (34863, Sigma-Aldrich), water, isopropyl ethanol, and water again before drying in an air stream. The slide was then further treated by a plasma cleaner (PDG-32G, Harrick Plasma) at 0.5 bar air pressure for 10 min at maximum power. A gasket (3 mm diameter, 1 mm thickness, GBL103280, Grace Bio-Labs) was carefully placed on the treated slide, and SLB buffer was immediately applied to all wells. Vesicles were then applied to the well to form SLB.

Residual vesicles were washed away with the SLB buffer by gently pipetting in the well without touching the membrane (as complete removal of all vesicles are rather impractical for SLB containing negatively charged lipid). Gag binding buffer was then introduced into the well. Sample protein solution or protein-RNA mixture was diluted or mixed in Gag binding buffer and incubated for 5 min before replacing the Gag binding buffer in the well, followed by recording of five 5-min movies at a frame rate of 100 Hz after 5 min incubation.

### Mass calibration curves and diffusion control

The contrast of a set of mass standards was measured for both the landing assay and MSPT on SLBs to convert the scattering contrast into molecular weight.

For landing assay mass standards, 50 nM bovine serum albumin (BSA) or 50 nM alkaline phosphatase was diluted in Gag binding buffer and equilibrated for 5 min at room temperature. The protein standard solution was added to the well of a landing assay set up and the landing events were recorded for 1 min.

For MSPT mass standards, POPC lipid vesicles containing 0.01 mol% biotinylated PE were used to form SLBs. 2.5 nM streptavidin was added to the SLB after the residual vesicles were washed away with Gag binding buffer. Unbound streptavidin was washed away with Gag binding buffer after 10 min incubation at room temperature, followed by the addition of 100 nM biotinylated BSA (A8549, Sigma-Aldrich), 100 nM biotinylated alkaline phosphatase (29339, Thermo Scientific), or 100 nM biotinylated Protein A (29989, Thermo Scientific). The unbound biotinylated protein was washed away after 5 min incubation at room temperature, and movies were recorded for 10 min for each protein. The composition of biotinylated PE and the concentration of streptavidin are the limiting factors to ensure there wasn’t too many particles in the FOV.

### Mass photometry data analysis

For a landing assay, background removal and contrast-to-mass conversion facilitated by standard curve were done using Discover^MP^ with default parameters.

For MSPT experiments, data processing was done using custom Python scripts. Background processing was done using a median background subtraction method. To detect particles laterally moving on the surface, this method calculates the pixel-wise temporal median image of a 1001-frame window (half window size n = 500) with the frame of interest in the middle and uses it as the static background to be removed for that particular movie frame.

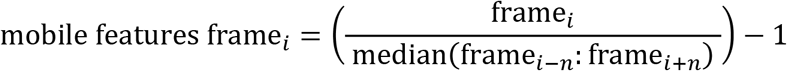

With this median background subtraction process, moving particles on the SLB would no longer appear to be distorted and can be properly fitted by point spread functions (PSFs) to extract contrast for each spot.

To identify spots in the background corrected movies, for each frame a Laplace filter was applied to filter out shot noise, followed by further thresholding to remove small promiscuous spots. A local maximum was identified for each of the candidate spots and its pixel coordinates were noted as the candidate pixel. A 13×13 pixels (84.4 nm per pixel) region of interest was marked for each candidate spot around the candidate pixel in the original image and fitted by the same PSF used in the Discover^MP^ software to extract particle contrast and location. Spots at very close distance were rejected at this step to guarantee good PSF fit.

After particle identification, trajectories were generated using the TrackPy package with minimum length filter set to 20 frames (200 ms), memory set to 1 frame, and maximum allowed step distance set to 5 pixels (422 nm).

The Python script used in background removal and particle identification was a combination of scripts from the Kukura Lab (University of Oxford) and the Schwille Lab (Max Planck Institute of Biochemistry) with specific modifications to suit the current needs.

For Gag-RNA complexes, the following considerations were taken into account: 1) as myr-Gag alone would not form complexes larger than Gag_3_ on SLB, it was assumed that no larger Gag-only complexes would form when RNA was added to the system; 2) under physiological conditions, HIV-1 would only package 2 copies of gRNAs into the newly assembled virions, thus it was assumed here the maximum RNA copy number in each complex species is 2; 3) when combined with mutated monomer gRNA 5’ UTR and incubated on the SLB, the largest complex was RNA_1_:Gag_4_ (data not shown), suggesting 2 RNA molecules need to exist and dimerize for more than 4 Gag proteins to be incorporated into the complex. All peak assignments regarding Gag-RNA complexes were made based on these three premises.

### Step detection algorithm and step size analysis

To detect mass change steps throughout trajectories, a C language implementation of the Kalafut-Visscher algorithm was used to perform denoising and step detection. The algorithm itself does not require any other input except for the time series itself. Previous studies have reported that the length of the time series and the position of potential change points can affect the step detection result and its accuracy. To address this issue, all individual trajectories from a certain group of experiments were concatenated first, and it was then divided into a series of *n* segments of equal length using a sliding window of 1000 frames. Each of these individual 1000-frame segments were used as input to the algorithm for analysis, avoiding bias from step positions and the length of the time series. With this sliding window segmentation procedure, step detection was repeated 1000 times shifting the start point by 1 frame in each iteration. After the step detection was done for every segment, the results were pooled together. For a particular time-spot in a trajectory, it had gone through 1000 iterations of step detection cycle, and the frequency of a returning result being a step was used to decide if it is a real change point identified at that particular location with a threshold of 0.75 (if a location returned more than 750 step detected results among the 1000 iteration, it is considered to be an actual step).

The sliding window length and the threshold are related to each other. A larger window length would result in less sensitive step detection, which can be compensated by choosing a smaller threshold. Considering the noise level of the system and the complexity of the Gag oligomerization process, a rather strict threshold was chosen in this project to avoid over-identification of steps.

## Acknowledgement

We would like to thank Silver Jõemetsa for providing help in optimizing SLB formation process. We would also like to thank Eric Foley from the Kukura Lab (University of Oxford) and Frederik Steiert from the Schwille Lab (Max Planck Institute of Biochemistry) for providing the original code for MSPT analysis. **Funding:** This work was supported by a grant from the NIH (U54AI170855) to J.R.W. and D.J.M. **Authors contributions:** A.X-Z.Z. designed and conducted the experiments, customized the code, analyzed the data and wrote the manuscript with input from all authors. J.A.H. made the original plasmids containing full-length Gag and NMT1 and optimized protein purification protocol. K.S. helped optimize the protein purification protocol and customize the code. D.P.M. provided insights into single molecule data interpretation. J.R.W. provided insights into data interpretation and helped conceptualize the assembly model. **Competing interests:** The authors declaring no competing interests. **Data and materials availability:** All data are available in the manuscript or the supplementary materials. Materials are available upon request.

