## Supplemental Figures for "Early HIV-1 Gag Assembly on Lipid Membrane with vRNA"

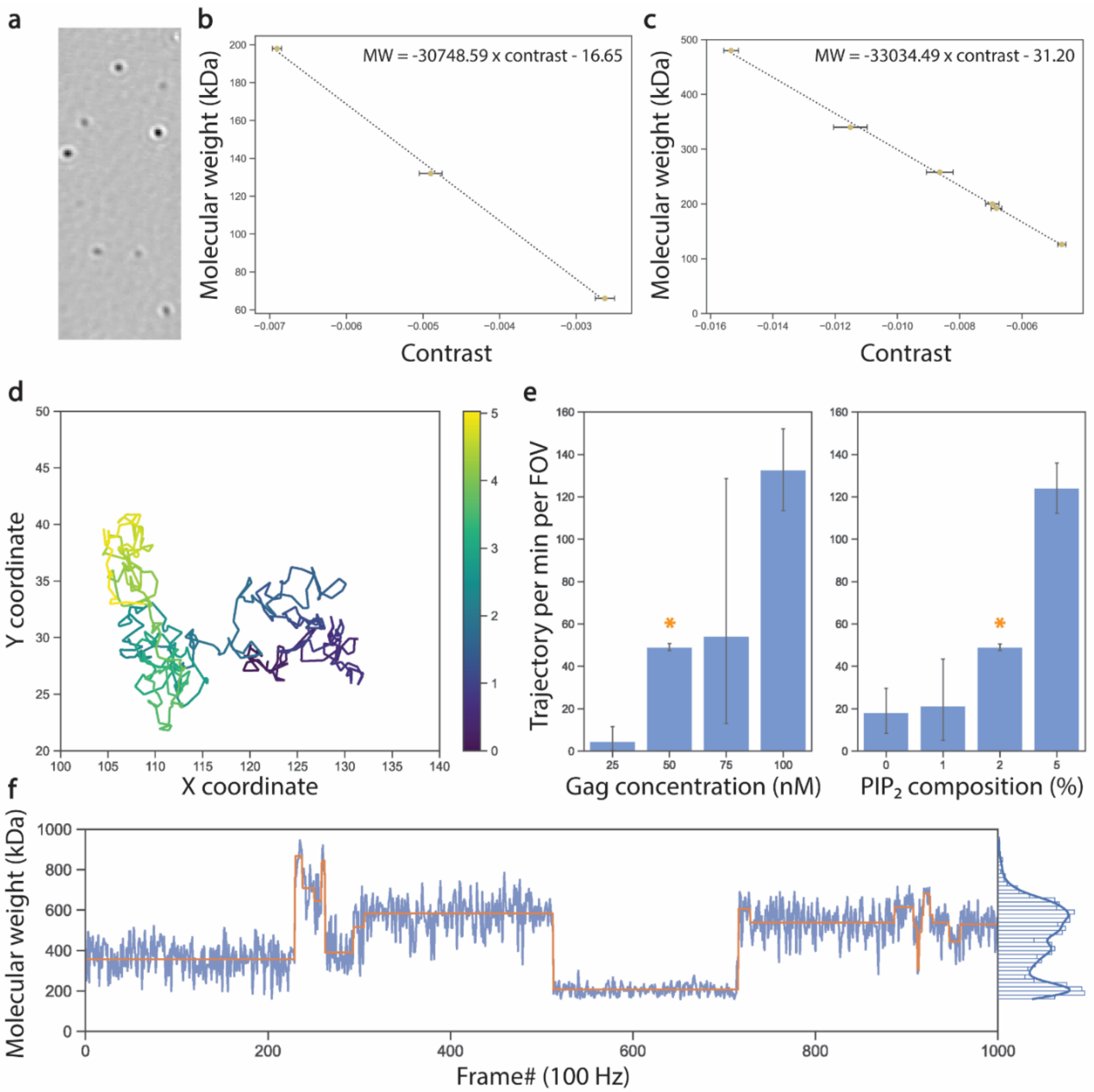


**Fig. S1** Samples of mass photometry data, optimization, and calibrations. (**a**) Typical contrast image from the mass photometer in an MSPT assay for myr-Gag diffusing on an SLB (POPC membrane with 2% PI(4,5)$P_2$). The FOV is 9 x 7 microns, composed of 150 x 60 pixels. (**b**) Sample calibration curve for landing assay. BSA was used as a standard protein with its monomer, dimer and trimer peaks being detected. (**c**) Sample calibration curve for MSPT assay. Bio-BSA and bio-AP were used as a standard protein on SLB containing 0.01% bio-PE preincubated with tetravalent streptavidin. (**d**) MSPT measurement of a single trajectory of one single particle over time, consisting of 500 frames collected over 5 s at a frame rate of 100 Hz. The trajectory is colored by time interval. (**e**) Average number of trajectories per FOV as a function of PI(4,5)$P_2$ composition at 50 nM myr-Gag (left), and as a function of myr-Gag concentration at 2% PI(4,5)$P_2$ (right). **(f)** Molecular weight of the particle in each frame over the trajectory, with raw data (blue) and the results of step-detection algorithm (orange) together shown in the plot. The histogram of molecular weight distribution is shown at the right.

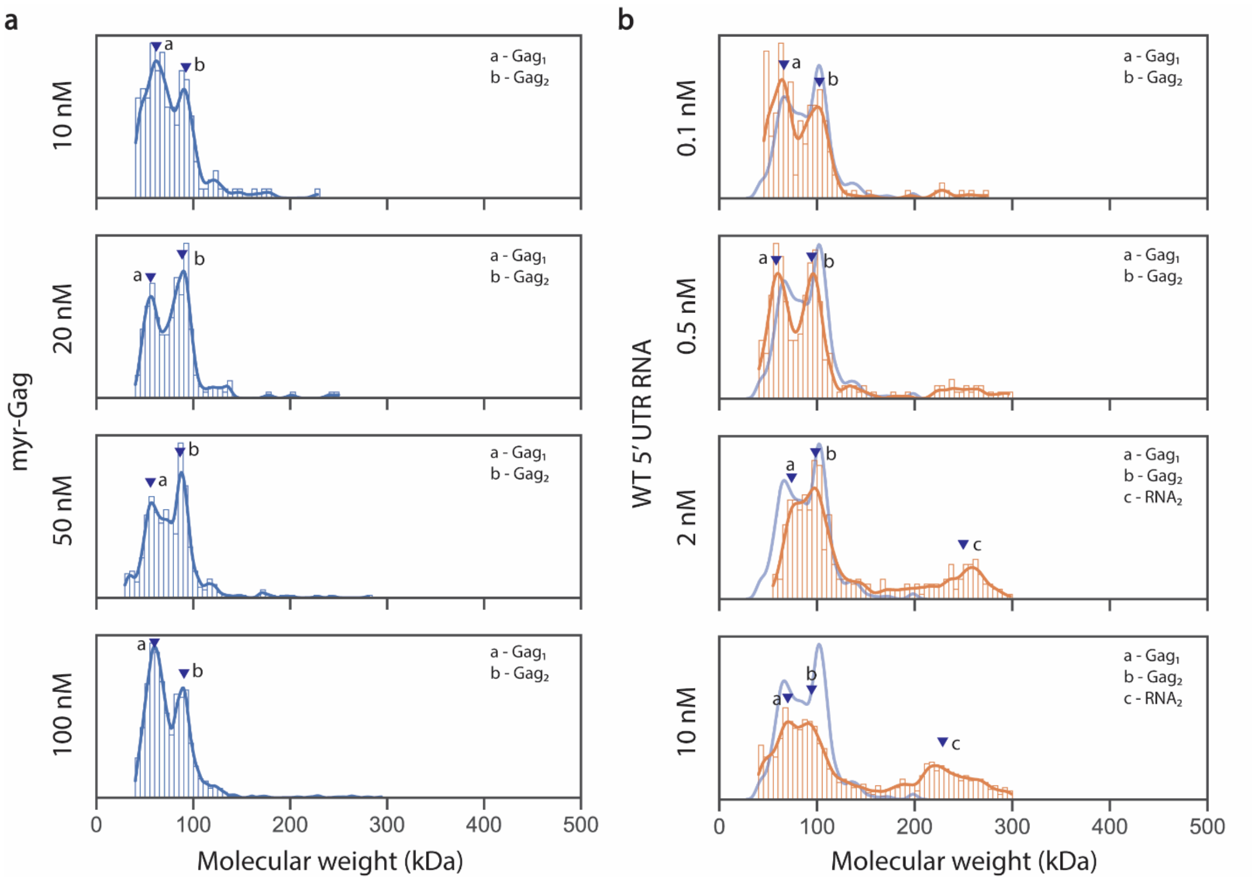


**Fig. S2** Molecular weight distribution of myr-Gag without and with WT 5' UTR RNA. **(a)** Molecular weight distribution probability density plot of 10 nM to 100 nM myr-Gag in binding buffer, measured by landing assay. **(b)** Molecular weight distribution probability density plot of 50 nM myr-Gag mixing with different concentrations of WT 5' UTR RNA in binding buffer, measured by landing assay. The light blue line indicates the molecular weight distribution of 50 nM myr-Gag in binding buffer. Labeled complex composition were estimated from kernel density plot peak molecular weights. All sample sizes are larger than 400 and collected from at least 3 different recordings.
